# Alizarin Red fluorescence imaging for nano calcification

**DOI:** 10.1101/2022.07.28.501840

**Authors:** Caterina Vanelli Coralli, Jing Xue, Adrian H Chester, Sergio Bertazzo

## Abstract

The formation of calcium phosphate (calcification) has been observed in a variety of healthy and diseased tissues and contributes to a wide range of pathologies. In physiological and pathological mechanisms, calcification begins at the nano scale and then develops into structures that range from a few micrometres to several centimetres. Fluorescence microscopy can be an efficient way to visualise such early calcification and its interaction with cells and proteins. The limited shelf life and high cost of commercial dyes for staining calcification, however, can be problematic when using this imaging method. Here, we aim to evaluate the effectiveness and stability of Alizarin Red (AR) as a fluorescent staining agent for nano and micro calcified structures. Our results show that AR staining for nano and micro calcified structures is a simple, reliable, effective, and quite inexpensive method to visualize calcification at the nano and micro scale in biological samples.

## Introduction

The formation of calcium phosphate (calcification) has been observed in a variety of healthy and diseased tissues and contributes to a wide range of pathologies, including cardiac disease^1-3^, cancer^4-7^, renal disease^8, 9^ and neurodegenerative disorders^10-13^. In physiological and pathological mechanisms, calcification begins at the nano scale and then develops into structures that range from a few micrometres to several centimetres.

Fluorescence microscopy can be an efficient way to visualise such early calcification and its interaction with cells and proteins. The limited shelf life and high cost of commercial dyes for staining calcification, however, can be problematic when using this imaging method.

Here, we aim to evaluate the effectiveness and stability of Alizarin Red (AR) as a fluorescent staining agent for nano and micro calcified structures. AR has been traditionally used to visualize calcification^14^ by naked eye or bright field microscopy^15-17^. We evaluate the effectiveness and stability of AR as a fluorescent staining agent for nano and micro calcified structures in aortas.

## Methods

### Samples

Aortic tissue samples from two patients were obtained from the Oxford Tissue Bank. The samples were fixed in formalin, embedded in paraffin, cut into 4 μm sections and mounted on glass slides.

### Sample staining

Before staining, the samples were deparaffinated and rehydrated using xylene, ethanol and Phosphate-Buffered Saline (PBS) washes.

Samples were stained for anti-alpha vascular muscle cell actin, DNA (DAPI), and calcium phosphate with OsteoSense 680EX (PerkinElmer NEV10020EX) or AR solution. The samples were permeabilised using a 0.1% Triton X-100 solution, then blocked in TBST with 5% BSA. Both samples were stained using rabbit polyclonal anti-alpha muscle cell actin I antibody (Abcam - ab5694) at a concentration of 1:5. The secondary antibody, Goat Anti-Rabbit IgG H&L (Alexa Fluor® 633, Thermofisher - SAB4600140), was applied at a concentration of 1:100.

AR solution was prepared by diluting 0.0625g of Alizarin Sulfonate (Merk - A5533-25G) powder in 100mL of distilled water. The pH was adjusted to near-neutral; the solution was then filtered and stored at 4ºC. The samples were incubated with AR in a dark environment for 10 minutes. The samples were then washed in PBS 8 times for 5 minutes each time. After staining with AR, the samples were washed in DAPI at a concentration of 1:20000.

For OsteoSense staining, the samples were t incubated with pure OsteoSense for 20 minutes. Two PBS washes, 10 minutes each were then carried out.

### Fluorescence microscopy

The samples were imaged on a Zeiss Airyscan 980_MP, using laser with excitation 488 nm and emission 590 nm for AR. Alex Fluor 633 and DAPI laser settings were used to image smooth muscle actin and DAPI, respectively.

## Results

Cardiac tissues often present nano and micro calcified spherical particles^18^, which make this type of tissue ideal for the evaluation of the capabilities of a fluorescent dye to image nano structures on calcified tissue. When stained with the commercial staining (OsteoSense) and imaged by confocal microscopy (Fig. 1a and 1b), aortas clearly present nanoparticles identifiable with considerable detail (Fig. 1b), and clearly the same structures seen in scanning electron micrographs^2^. Images of similar quality are obtained using AR solution (Fig. 1c and 1d) displaying the spherical particles with the same structure and frequency as observed when using commercial staining.

**Figure 1:**
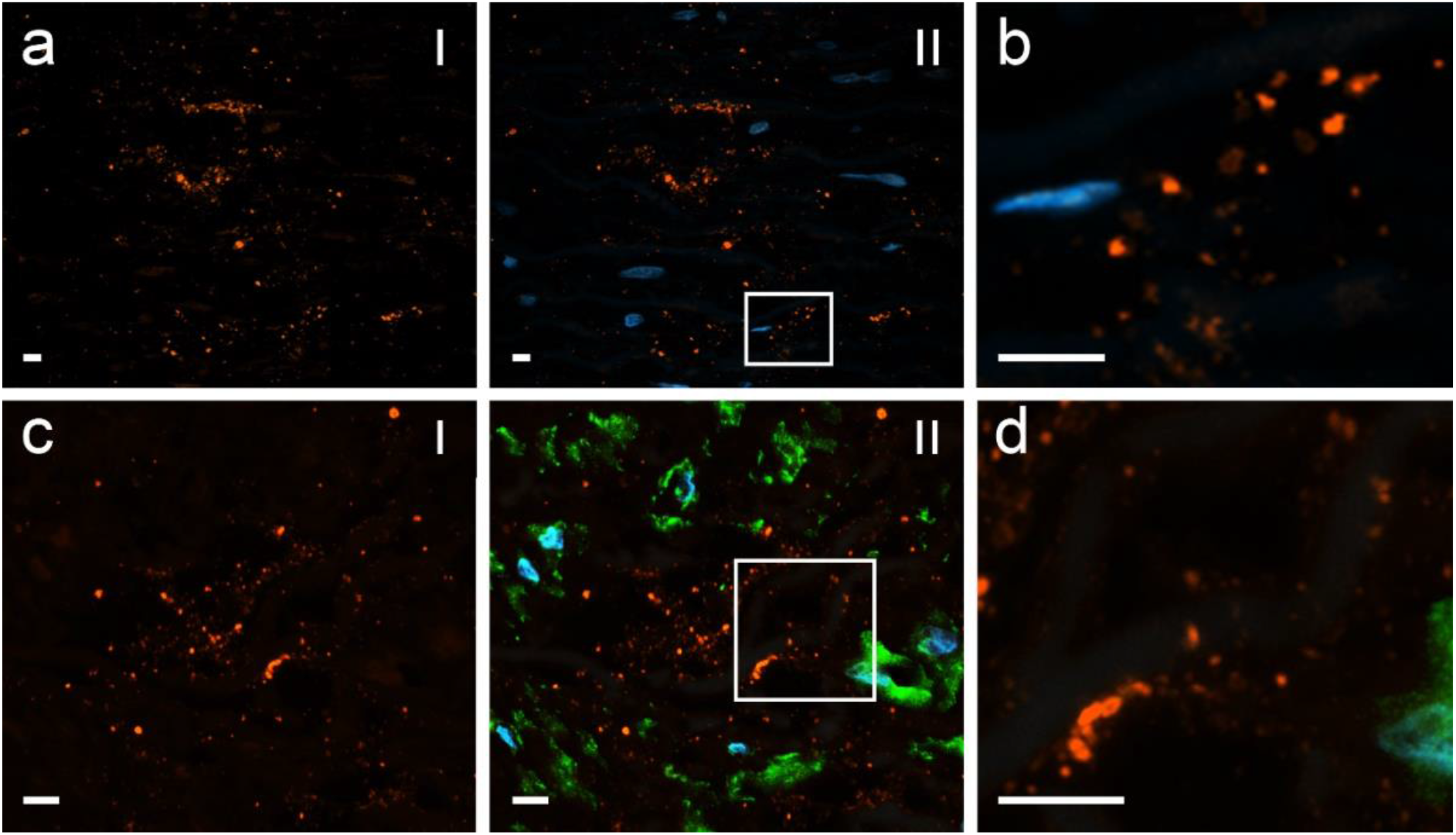
Calcified human aorta stained with Osteosense and AR. *a* Aorta stained with Osteosense (orange - *I*) and Dapi (blue - *II*). *b* Higher magnification of region highlighted by square in *aII. c* Aorta stained with AR solution (orange - *I*), vascular smooth muscle cells actin (green) and DAPI (blue - *II*). *d* Higher magnification of region highlighted by square in *cII*. Scale bar: 5 μm.

The compatibility of AR with immunostaining techniques was assessed by imaging tissue stained with an anti-alpha muscle cell actin I antibody (Fig. 1c and 1d). No cross-reactivity between the antibody and AR was observed, suggesting that AR staining can be successfully implemented in immunostaining protocols.

One of the most common concerns related to the use of AR for staining calcified structures that will be imaged by fluorescence microscopy, is that anecdotally it is believed that AR staining is not stable. To evaluate the stability of AR staining, samples were imaged at 24 hours, then 5, 10, and 15 days after staining. The micrographs (Fig. 2) show no change in the staining of the spherical particles. Moreover, it is clear from the images that no staining is present in any other structure even after 15 days (Fig. 2d), demonstrating that there is no time-related contamination of AR to other structures in the sample.

**Figure 2:**
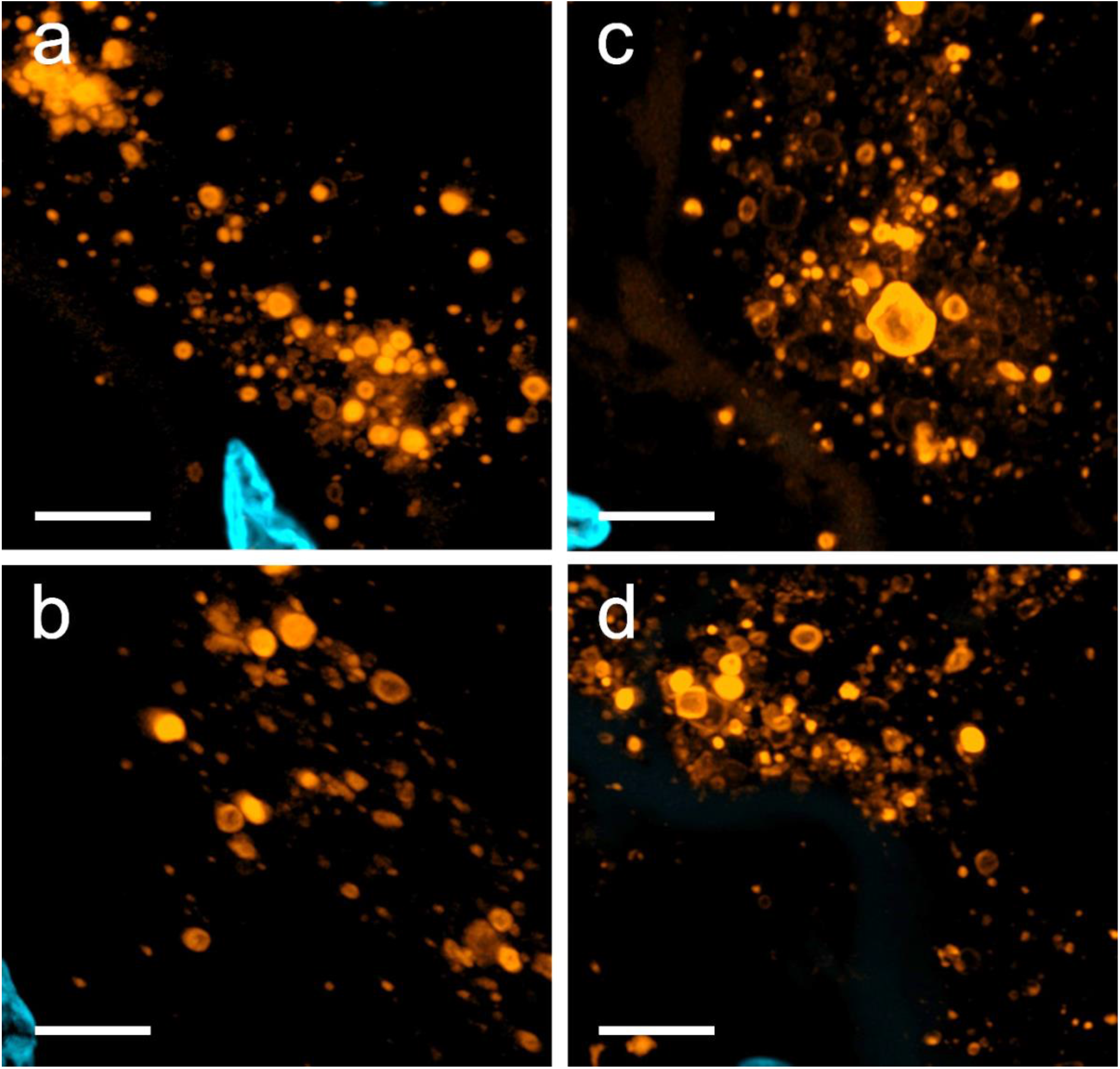
Calcified human aorta at different times after staining with AR. The tissues stained with AR and DAPI were imaged using confocal microscopy at various times after staining. Calcified microparticles are coloured in orange while cell nuclei are displayed in blue. *a* 24 hours post-staining. *b* 5 days post-staining. *c* 10 days post-staining. *d* 15 days post-staining. Scale bar: 5 μm.

## Conclusion

Considering the results obtained, it is clear that AR staining for nano and micro calcified structures is at least equally efficient as commercial staining agents used for the same purpose. Perhaps more importantly than efficiency is the stability of staining with AR. We have tested the reproducibility and confidence of experiments: even if non-specific staining might occasionally occur, we have not found any lack of specificity when using AR in calcified structures.

Moreover, AR solutions are stable at 4°C over months. This is in strong contrast to the commercial staining, which even if stored at -80°C will degrade after a couple months and will start to stain other structures in the tissue.

Taking all our findings together, it is clear that AR staining for nano and micro calcified structures is a simple, reliable, effective, and quite inexpensive method to visualize calcification at the nano and micro scale in biological samples. In particular, the ability to be used together with regular immunostaining, combined with the fact that AR staining does not degrade (and therefore, does not stain other, non-calcified structures in the sample), can contribute considerably to the study of cardiovascular calcification and other pathological calcification present in soft tissues.

